# Interferon-β paracrine signaling mediates synergy between TLR3 and TRIF-independent TLR pathways

**DOI:** 10.1101/2025.05.13.653801

**Authors:** Erick I. Salvador Rocha, Kayla Samo, Kathryn Miller-Jensen

## Abstract

Cytokine dysregulation during microbial infections can dangerously increase the severity of associated diseases. Non-additive responses to simultaneous activation of toll-like receptor (TLR) signaling pathways are one potential contributor. Here we explored mechanisms underlying the TLR2/MyD88 and TLR3/TRIF-mediated responses in macrophages for canonical pro-inflammatory and antiviral targets including TNF, IL-6, IL-12p40, IFN-β and CXCL10. We found that all targets exhibited characteristic levels of synergistic cytokine activation that varied with TLR-ligand concentration and exposure time. These trends were conserved when the TLR3-ligand polyI:C (PIC) was combined with either of the TLR2-mediated pathogens *S. aureus* or *L. pneumophila*. Using pharmacological inhibitors, genetic knockouts, and recombinant cytokines, we explored how TLR2-TLR3-induced paracrine signaling via TNF and IFN-β affected synergy. While TNF synergy is paracrine-independent, IFN-β contributes to synergistic activation of IL-6 and IL-12p40. When IFN-β was directly combined with P3C or *S. aureus* infection, low concentrations produced modest synergy for both, while high concentrations increased IL-6 but antagonized IL-12p40. Thus, multiple mechanisms regulate TLR2/TLR3-mediated synergistic cytokine activation in macrophages, including type I interferon modulation of TRIF-independent pathogen stimulation that increases inflammatory signaling. These findings have important implications for therapeutic modification of cytokine-driven inflammation, including cytokine storms and the development of vaccine adjuvants.

## Introduction

The ubiquitous nature of microbes and their range of ecological relationships with mammalian hosts–including commensalism, symbiosis, and parasitism–is a discrimination challenge for the immune system, because simultaneous complex signals need to be detected and processed. Innate immune cells use a limited repertoire of pattern-recognition receptors (PRRs) to recognize conserved structures called pathogen-associated molecular patterns (PAMPs). The Toll-like receptors (TLRs) are one well-characterized group of PRRs that are present in many innate immune cells (Takeuchi & Akira, 2010). Macrophages are among the first cells to detect and respond to the activation of their TLRs by producing cytokines that engage other immune cells and mount an appropriate inflammatory response (Arango Duque & Descoteaux, 2014). The individual TLR-ligand interactions and their corresponding signaling pathways have been described extensively in macrophages (Alexopoulou et al., 2001; Hemmi et al., 2002; Horng et al., 2002; Kagan et al., 2008; Kawai et al., 1999; Takeuchi et al., 2001; Yamamoto et al., 2003). However, the combinatorial nature of TLR-signaling in response to multiple microbial inputs and in the context of co-infections is an important area of research.

Upon activation, TLRs recruit and signal through two primary signaling adaptors, MyD88 and TRIF, resulting in the activation and downstream signaling of key transcription factors necessary to produce inflammatory cytokines (Kawai & Akira, 2006). Simultaneous TLR-pathway activation can lead to both synergistic and antagonistic inflammatory responses, where synergy is influenced by the crosstalk between the MyD88- and TRIF-signaling pathways regardless of the specific TLR being activated or its cellular localization (Bagchi et al., 2007; Napolitani et al., 2005; Sato et al., 2000; Thaiss et al., 2016). Among TLRs, TLR3 is often studied, as it is activated at the endosomal level, recognizes viral double-stranded RNA, and, most importantly, is the only TLR that signals solely through the TRIF-signaling pathway. On the other hand, TLR2 is activated at the surface level of innate immune cells, can recognize a wide range of microbial inputs from parasitic, bacterial, fungal, and viral origin, and signals exclusively through the MyD88 pathway (Fitzgerald & Kagan, 2020).

When exploring non-additive responses with TLR2 and TLR3, the cytokines interleukin 6 (IL-6) and IL-12p40 have been consistently classified as synergistic, while the inflammatory cytokine tumor necrosis factor (TNF) has been reported to be synergistic in some contexts but not others (Ilievski & Hirsch, 2010; Lin et al., 2017; Liu et al., 2015; Suet Ting Tan et al., 2013). Moreover, many studies use a single dose of TLR ligands and a fixed duration of ligand exposure; research on dose-dependent variation in synergy has been limited, even though non-additive TLR activation can be a function of both dose and time. Given that immune cells encounter different quantities of PAMPs, understanding how TLR-ligand dose impacts their response is important.

TLR-mediated responses are further regulated by paracrine signaling, in which cytokines secreted by cells can influence the activation state and response of neighboring cells. In the context of the ligand lipopolysaccharide (LPS), which activates both the MyD88 and TRIF-signaling pathways through TLR4, TNF has been implicated as a positive feedback amplifier in inflammatory responses, while type I interferons can modulate both pro-inflammatory and antiviral pathways (Alexander et al., 2021; Gottschalk et al., 2019; Hu et al., 2008; Lee et al., 2009; Song et al., 2015). Although it has been shown that cell-cell communication through TNF and IFN-β paracrine signaling can amplify TLR-mediated inflammatory responses (Shalek et al., 2014; Xue et al., 2015), a role for these cytokines in mediating synergy between combinations of TLRs has not been widely explored.

In this study, we sought to elucidate the mechanisms underlying cytokine synergy in the TLR2-TLR3 combinatorial response. We characterized the synergistic behaviors of canonical pro-inflammatory and antiviral targets (i.e., TNF, IL-6, IL-12p40, IFN-β, and CXCL10) and found that synergistic activation varied in relation to time and TLR-ligand dose. We demonstrated these trends are largely conserved in more biologically complex stimulations with the bacteria *S. aureus* and *L. pneumophila* in combination with a TLR3-ligand. We examined the contributions of the TNFR- and IFNAR-signaling pathways using a combination of pharmacological inhibitors, genetic knockouts, and recombinant cytokines, and found a role for TLR3-induced IFN-β in modulating TLR2-TLR3 synergistic activation of IL-6 and IL-12p40. Finally, we demonstrated that, when combined with a TLRs that signal via MyD88, IFN-β was sufficient to synergistically amplify IL-6 and antagonize IL-12p40 secretion to varying extents across all TLRs tested. Our findings reveal dose-dependent modulation of the TLR2-TLR3 synergistic response and distinct regulatory mechanisms for TNF, IL-6, and IL-12p40 synergy, with IFN-β directly influencing the non-additive responses of IL-6 and IL-12p40 in response to TLR3 and MyD88-dependent ligands.

## Results

### Non-additive responses exhibit different trends between transcription and secretion that vary across cytokines

To study how TLR2 and TLR3 responses combine during the early phase of the inflammatory response, we stimulated bone-marrow derived macrophages (BMDMs) with the TLR2 ligand Pam3CSK4 (P3C), the TLR3 ligand poly(I:C) (PIC), or both (dual stimulation) (Fig. 1A). We selected four key cytokines involved in the response: TNF and IFN-β, which are primary paracrine signals downstream of TLR2 and TLR3, respectively (Fitzgerald & Kagan, 2020; Siebeler et al., 2023); and IL-6 and CXCL10, which are secondary response signals requiring translation of intermediate proteins for their activation (Tong et al., 2016). We measured transcription using quantitative RT-PCR over a time course up to 8 hours, and protein secretion by ELISA over 24 hours in response to P3C, PIC, or dual stimulation. The degree of non-additivity in the response at each time point was calculated as (response to P3C+PIC)/(response to P3C + response to PIC), with values > 1 indicating synergy (Fig. 1B). This definition is consistent with previous studies (Suet Ting Tan et al., 2013).

**Fig. 1.**
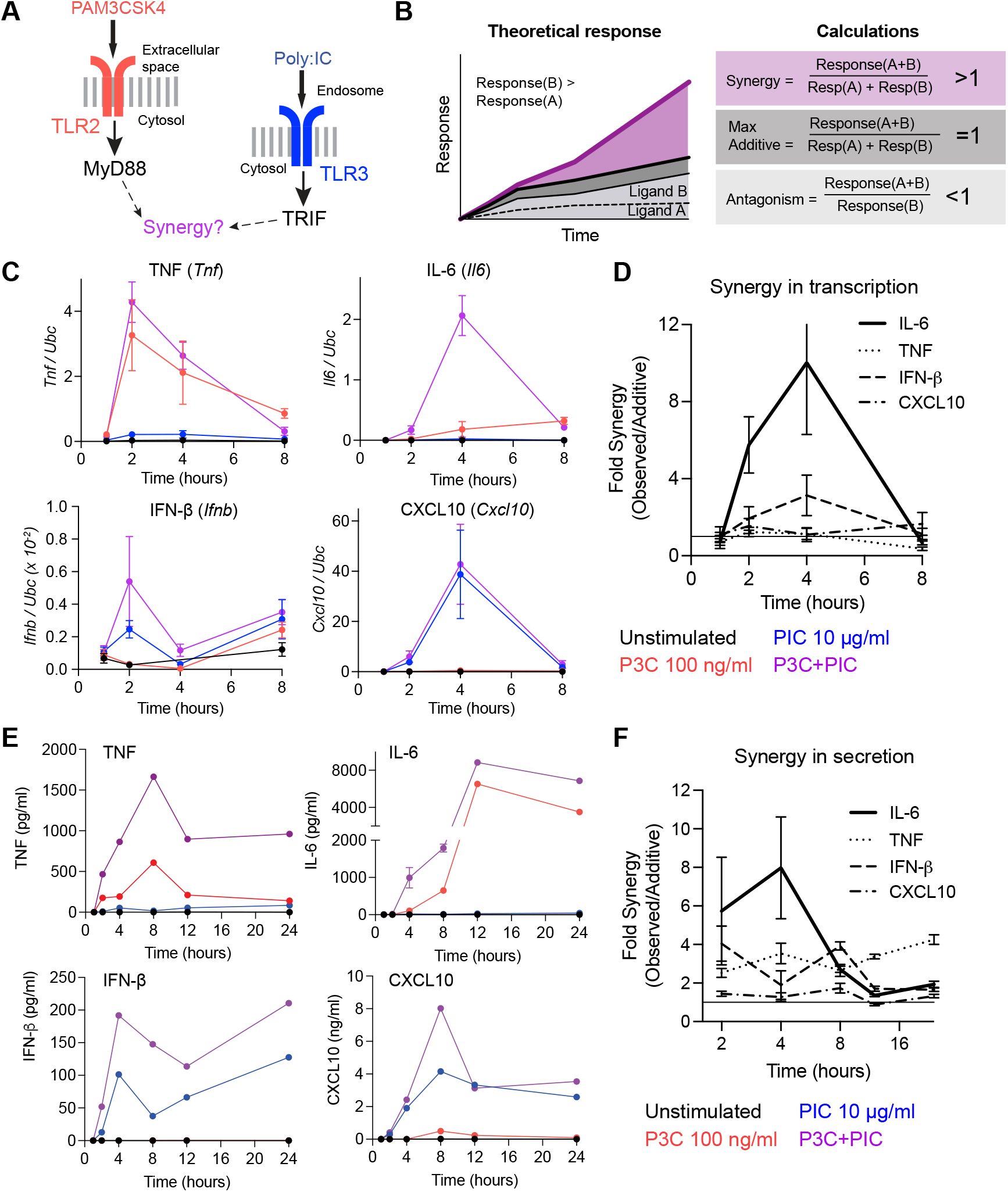
Synergy between TLR2 and TLR3 ligands exhibits different trends between transcription and secretion that vary across cytokines. (A) Schematic of TLR-signaling pathways. (B) Schematic defining non-additive responses in a time course response graph and the associated calculations. (C) RT-qPCR measurements of indicated genes in BMDMs stimulated with the indicated doses of Pam3CSK4 (P3C) and/or poly(I:C) (PIC) at 1, 2, 4, and 8 hours. mRNA levels for each target are normalized to the housekeeping gene Ubc (n = 3). (D) Calculated fold synergy of the P3C+PIC transcriptional response for all measured genes across timepoints. (E) ELISA measurements of protein concentrations from 1-24 hours for the conditions in (C). (F) Calculated fold synergy (as described in D) of the P3C+PIC secretion response for all measured cytokines across timepoints. Data are presented as mean SD of n = 3 biological replicates.

We first examined synergy in transcription of the four targets of interest. Transcription of all targets peaked between 2-4 hours and declined almost back to baseline by 8 hours (Fig. 1C). *Tnf*, which is activated strongly by P3C alone but weakly by PIC alone, did not exhibit synergy at the transcript level. In contrast, *Il6*, which is also more strongly activated by P3C than PIC alone, showed a more than 5-fold increase in peak transcription relative to the additive response when PIC was combined with P3C (Fig. 1D). The PIC gene targets *Ifnb* and *Cxcl10* also showed distinct behavior: *Ifnb* exhibited synergy in transcription, but *Cxcl10* did not (Fig. 1C-D). Altogether we found that TLR2-TLR3 synergy in transcription varied by target.

We next examined synergy in protein secretion for the same four targets. Similar to the transcriptional dynamics, secretion of all targets peaked between 8 and 12 hours and, with the exception of IFN-β, declined or plateaued by 24 hours (Fig. 1E). All targets exhibited significant synergistic activation, although to varying degrees (Fig. 1F). IL-6 exhibited the highest synergy at 4 hours, consistent with the transcription results. TNF exhibited 3-4-fold synergy that stayed relative constant across time. The difference we observed in TNF synergy for transcription versus secretion is consistent with conflicting prior studies that reported TLR2-TLR3 synergy for TNF secretion (Bagchi et al., 2007; Ilievski & Hirsch, 2010), but not significantly for TNF transcription (Lin et al., 2017). This result is also consistent with a role for TRIF signaling in stimulating TNF translation and secretion (Caldwell et al., 2014; Gais et al., 2010).

To determine if synergy is indeed occurring due to increased translation and not just increased secretion, we blocked protein secretion by treating cells with Golgi Plug™, a Brefeldin A (BFA)-containing compound, during stimulation. We then lysed the cells and measured the intracellular concentrations of TNF, IL-6, IFN-β, and CXCL10 by ELISA. For TNF, IL-6, and IFN-β, we observed a synergistic increase in protein level, demonstrating that production is being amplified at the level of translation, although the synergy for IL-6 protein expression was reduced compared to the protein levels in supernatants (Fig. S1A). In contrast, CXCL10 protein concentration in the lysates fell under the range of the ELISA assay (62.5 pg/ml), consistent with its classification as an interferon-stimulated gene (ISG) and its requirement for secretion of IFNs. Overall, our data suggest that distinct regulatory mechanisms mediate the synergy of different pro-inflammatory and antiviral targets.

To analyze synergy in TLR2-TLR3-stimulated secretion more globally, we collected supernatants at 4 and 8 hours and performed a multiplex bead-based immunoassay of 32 cytokines and chemokines (C/Cs). For the 21 C/Cs significantly activated by TLR2 and/or TLR3, we performed hierarchical clustering analysis on the overall secretion data. We found that they clustered based on the individual ligand activation and dynamics: targets activated by P3C (TLR2) or PIC (TLR3) stimulation clustered together, while two other clusters contained targets activated by by 4h and 8h (Fig. S1B). We then calculated values of synergy at 4h and 8h and re-clustered the C/Cs (Fig. S1C). Clustering based on synergy revealed different groups of C/Cs. Cluster 1 targets, including CCL2 and IL-1β, showed no synergy or, in some cases, antagonism. Cluster 2, which contained CXCL10 as well as IL-10, exhibited low synergy at 4 hours but no synergy or antagonism by 8 hours. Cluster 3, which contained TNF and CXCL1, exhibited low synergy at both time points. Finally, Cluster 4, which included IL-6 and IL-12p40, exhibited the strongest synergy at 4h. We confirmed IL-12p40 synergy by ELISA (Fig. S1D). Overall, the chosen targets appear to span the range of synergy patterns observed for inflammatory cytokines and chemokines.

### Dose matrix reveals minimum thresholds and dose-dependent variation in TLR2-TLR3 synergy

We next considered how the dose of P3C and PIC would affect TLR2-TLR3 synergy in protein secretion. We focused on the 4-hour time point because it exhibited the highest level of synergy in secretion across targets. Previous studies have demonstrated dose-dependent variation in cytokine secretion in the context of TLR2-TLR4 and TLR3-TLR7 combinations (Gottschalk et al., 2019; Suet Ting Tan et al., 2013). However, the TLR2-TLR3 synergy in secretion has not been well elucidated. To understand the dose-dependence of non-additivity in the TLR2-TLR3 response, we stimulated BMDMs with increasing doses of P3C or PIC and all the pairwise combinations. We collected the supernatants at four hours, and measured the protein secretion of TNF, IL-6, IL-12p40, IFN-β, and CXCL10 by ELISA and calculated fold synergies for all pairwise combinations.

We first considered how PIC amplified expression of P3C-mediated activation of TNF, IL-6, and IL-12p40. We observed a secretion threshold for all three cytokines that depends on the dose of the TLR2 ligand P3C, with robust cytokine secretion at doses above 10 ng/ml (Fig. 2A). Interestingly, the addition of PIC did not lower the P3C dose threshold but rather resulted in an increase in the area under the curve, indicating that PIC magnifies the pro-inflammatory response strength rather than reducing the TLR2 activation threshold. There was little difference in the amplification observed at different doses of PIC. For all dose combinations, peak secretion for TNF and IL-12p40 occurred at the 100 ng/ml P3C dose, with TNF secretion plateauing and IL-12p40 secretion decreasing at 1000 ng/ml. This observation supports possible anti-inflammatory regulation at higher concentrations which could be mediated by IL-10, as has been observed in TLR2-TLR3 stimulations in human dendritic cells (DCs) (Re & Strominger, 2004). By comparison, IL-6 secretion continued to increase in a dose-dependent manner after crossing the 10 ng/ml threshold.

**Fig. 2.**
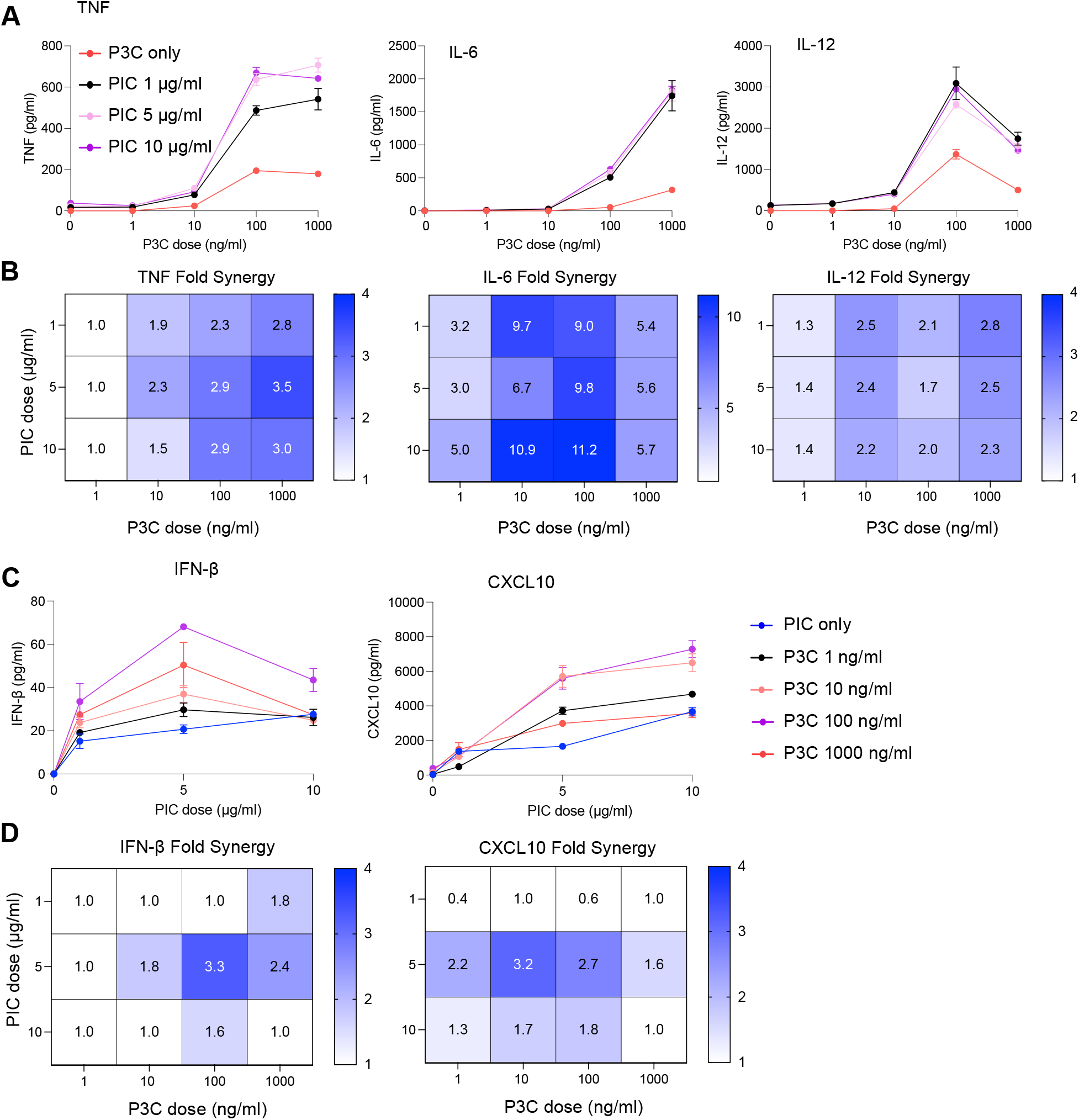
TLR2 and TLR3 ligands exhibit minimum thresholds and dose-dependent variation in TLR2-TLR3 synergistic activation of cytokines. BMDMs were stimulated with increasing doses of P3C (0, 1, 10, 100, 1000 ng/ml) in combination with increasing doses of PIC (0, 1, 5, 10 ug/ml). (A-B) Secretion (A) and fold synergy calculations (B) of pro-inflammatory cytokines TNF, IL-6, and IL-12p40. (C-D) Secretion (C) and fold synergy calculations (D) of antiviral cytokines IFN-β and CXLC10. Data are presented as mean SD of n = 2 biological replicates.

Synergistic activation of TLR2 stimulation by TLR3 for each cytokine was consistent across doses. PIC induced an approximately 3-fold synergistic increase in TNF at 100 and 1000 ng/ml (Fig. 2B). PIC induced an approximately 6-9-fold synergistic increase in IL-6 at P3C doses > 10 ng/ml, although overall secretion of IL-6 was very low at 10 ng/ml P3C. The calculation of synergy across the titration matrix reveals that, like overall IL-12p40 secretion, the highest levels of synergy occur at the lowest PIC dose of 1 µg/ml and at either 10 or 1000 ng/ml of P3C, which is different than its maximum secretion dose of 100 ng/ml. Such differences reveal different regulatory mechanisms of synergy than those of IL-6.

We next examined how P3C amplified PIC-mediated activation of the antiviral targets IFN-β and CXCL10. We did not observe the same thresholding behaviors as TNF and IL-6, possibly due to the narrower PIC dose range (Fig. 2C). The secretion peaks for these cytokines differed, with CXCL10 peaking at the highest PIC dose (10 µg/ml) and IFN-β peaking at a medium PIC dose (5 µg/ml). Despite these differences, P3C increased secretion of both targets in a dose-dependent manner (Fig. 2C). The synergy for these antiviral targets also exhibited less pronounced thresholding, with the highest levels of synergy occurring in the mid-range of the titration matrix rather than at the dose extremes (Fig. 2D). Specifically, the 5 µg/ml PIC dose yielded the strongest synergy for both antiviral targets.

Since TNF secretion and synergy were not substantially reduced with BFA (Fig. S1A), we next explored if TNF synergy results from P3C+PIC activating a greater fraction of cells or by stimulating higher cytokine production per cell. BMDMs were stimulated with either a moderate (30 ng/ml) or high (100 ng/ml) dose of P3C combined with 10 µg/mL PIC in the presence of Golgi Plug™, and TNF expression was measured after four hours by flow cytometry. At 30 ng/ml P3C, adding PIC doubled the fraction of TNF+ cells from approximately 30% to 60%, and it tripled the fraction of IL-12p40+ cells from around 4% to 12% (Fig. S2A-B). At 100 ng/ml P3C, the TNF+ fraction of cells approached 60%, but the addition of PIC did not significantly increase this fraction. However, at this P3C dose the IL-12p40+ fraction had a significant but small increase with the addition of PIC. Moreover, regardless of the P3C dose, the dual stimulations showed no significant difference between the TNF+ or the IL-12p40+ fractions, suggesting that these populations might reach a saturating percent of activated cells (Fig. S2B). Additionally, for both P3C doses, PIC significantly increased the mean fluorescence intensity of TNF+ and IL-12p40+ cells at both P3C doses, indicating higher per-cell production of TNF (Fig. S2C). When we normalize the overall TNF and IL-12p40 secretion from our dose titration matrix (Fig. 2) to the fraction of positive cells for these cytokines, we observe that there are similar levels of cytokine secretion per cell for the single ligands and the moderate P3C dual stimulation, but that there is a much larger increase in the per cell secretion of TNF and IL-12p40 at the high P3C dual stimulation (Fig. S2C) These results suggest that at lower P3C doses, synergy with PIC results from more cells crossing the TNF and IL-12p40 activation threshold, whereas at higher P3C doses, synergy primarily reflects enhanced per-cell cytokine production. Given the negative impact of BFA on IL-6 synergy, it was not possible to explore if this trend held for IL-6.

### Poly(I:C) induces a synergistic inflammatory response in combination with *S. aureus* and *L. pneumophila*

Our results suggest that the TLR3-mediated antiviral response that signals through TRIF can synergize with TLR2 activation, which signals via MyD88, to increase the proinflammatory response. We next considered whether TLR3 activity would also synergize with TLR7/8 activation, another TLR receptor that signals through MyD88 independent of TRIF (Fig. S3A). We stimulated BMDMs with 100 ng/ml R848, 10 μg/ml PIC, or the combination, and measured protein secretion by ELISA at 2, 4, and 8 hours. We found that PIC combined with R848 to increase secretion of all measured targets (Fig. S3B). At least a 2-fold synergy was observed for TNF, IFN-β, and IL-6 (Fig. S3C). Although the secretion levels were more modest than those observed with P3C/TLR2, the patterns of synergy observed for TLR7 were very similar to those observed for TLR2.

To explore if the same trends for combined activation of TLR3 and TRIF-independent TLRs would be observed in a more biologically relevant context, we combined PIC stimulation with bacteria that produce a strong TLR2-mediated inflammatory response, specifically *Staphylococcus aureus* and a replication-deficient *Legionella pneumophila* (Fig. 3A). *S. aureus* is a commensal and sometimes an opportunistic Gram-positive bacterium that possesses lipoteichoic acids (LTA) and peptidoglycan which are recognized by TLR2 (Askarian et al., 2018). *L. pneumophila*, an opportunistic Gram-negative bacterium, possesses lipoproteins and flagellin that activate TLR2 and TLR5, respectively, as well as a poorly recognized LPS that can weakly activate TLR4 (Liu & Shin, 2019). However, we used a mutant *L. pneumophila* strain that does not possess flagellin or a functional secretion system (*ΔflaAΔdotA)* in order to focus on TLR2 recognition (Ren et al., 2006). We infected BMDMs with these two bacteria at a MOI of 10 in combination with a dose of 10 µg/ml PIC. We measured cytokine secretion of TNF, IL-6, IL-12p40, IFN-β, and CXCL10 over a time course of 2, 4, and 8 hours, and then calculated the degree of non-additivity as we previously described (Fig. 1B).

**Fig. 3.**
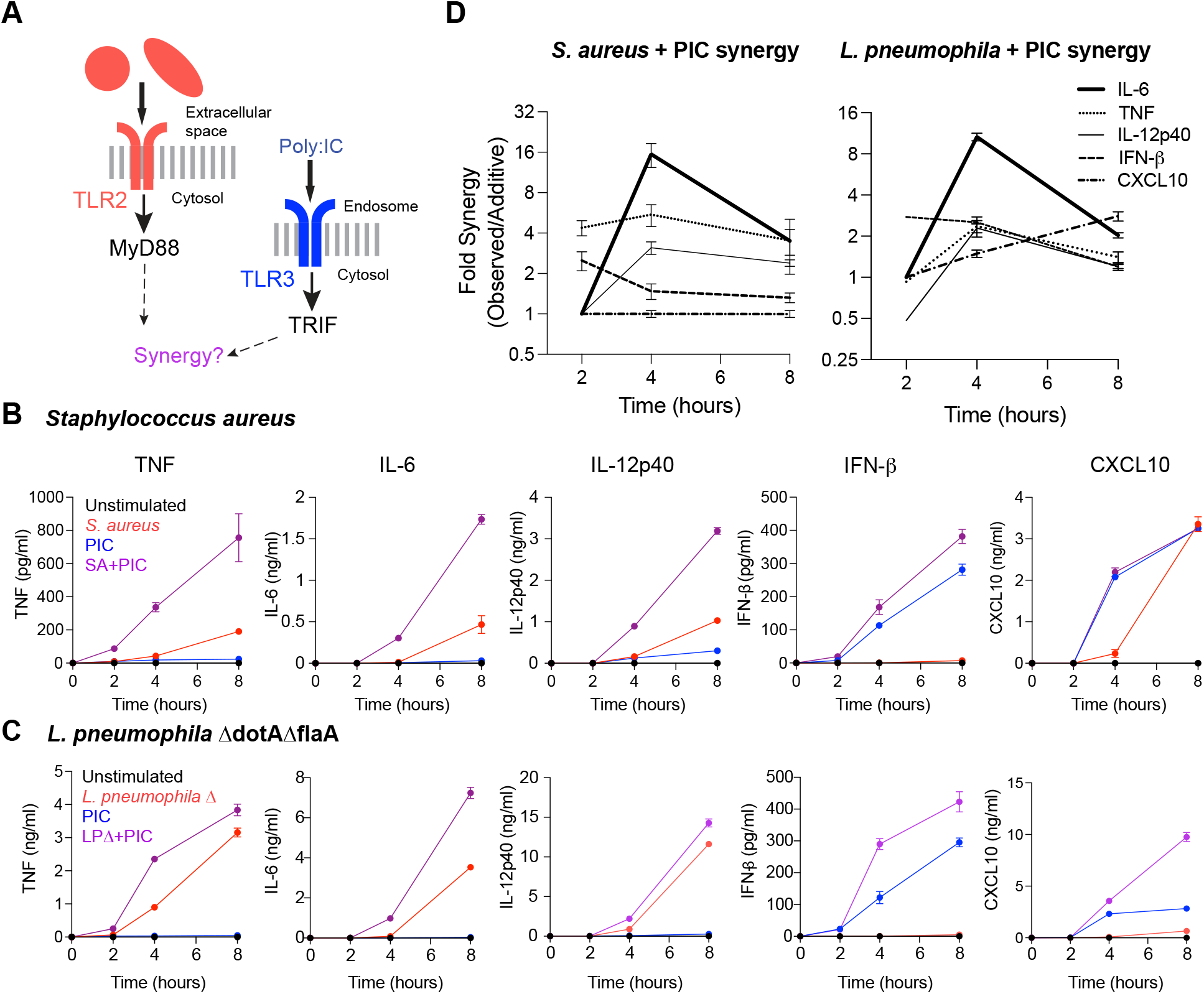
Poly(I:C) stimulation of TLR3 combines with TLR2 stimulation in the context of S. aureus and L. pneumophila infection to produce a synergistic inflammatory response. (A) Schematic of stimulation combinations. (B-C) BMDMs were infected with S. aureus (B) or mutant L. pneumophila strain Δ flaAΔdotA (C) at an MOI = 10 alone or in combination with poly(I:C). TNF, IL-6, IL-12p40, IFN-β, and CXCL10 secretion was measured in the supernatants by ELISA at 2, 4, and 8 hr post-infection. (D) Fold Synergy calculations of the observed dual response compared to the maximum expected additive response across timepoints. Mean and SD error bars of n = 2 biological replicates.

*S. aureus* infection alone produces moderate levels of TNF, IL-6, and IL-12p40 secretion that have a continuous increase between the 4- and the 8-hour timepoints (Fig. 3B). Combining this bacterium with PIC caused higher levels of cytokine secretion, as evidenced by the larger area under the curve. By comparison, *S. aureus* infection alone does not drive a strong IFN-β response as the secretion levels fall under the range of detection of the ELISA assay (12.5 pg/ml), and the combination with PIC causes a moderate increase of the area under the curve when compared to PIC alone (Fig. 3B). CXCL10 secretion in response to *S. aureus* exhibits a quick rise between the 4- and 8-hour timepoints, which is interesting given that this infection alone does not result in strong IFN-β secretion, suggesting possible activation of other IFN pathways. The combination of *S. aureus* with PIC does not result in synergy of CXCL10 at any timepoint, thus indicating no crosstalk.

*L. pneumophila ΔflaAΔdotA* infection alone produces higher levels of TNF, IL-6, and IL-12p40 secretion than those observed in our *S. aureus* model of infection, and their increase is observed at the 2-hour timepoint and continues to rise for the 8 hours of our assay. While the addition of PIC to the *Legionella* infection increases the secretion of TNF, IL-6, and IL-12p40, the overall increases to the area under the curve are more modest than those of *S. aureus* + PIC. Similar to *S. aureus* infection, *Legionella* infection alone does not drive a strong IFN-β response, as the secretion levels also fall below the range of the ELISA assay, and the combination with PIC also causes a moderate increase of the area under the curve when compared to PIC alone. However, the behavior of CXCL10 secretion is different between these two models because *Legionella* infection does not show high secretion of CXCL10 but in combination with PIC causes a synergistic effect (Fig. 3C).

Overall, when comparing the trends between stimulation with the TLR2 ligand P3C (Fig. 1E-F) and our two models of TLR2-mediated bacterial inflammation, we observe several consistent trends. These include an overall peak in synergy at 4 hours for most targets, and similar fold synergy increases across targets, with IL-6 consistently displaying the highest increase (>10 fold; compare Fig. 1E and Fig. 3D). This is also true for TLR7/8-TLR3 synergy (Fig. S3). Although there is some variation in the extent and timing of synergy, suggesting some fine tuning across models, the general mechanism mediating synergy, especially for IL-6, appears to be conserved.

### TLR2-TLR3 synergy of IL-6 exhibits type I IFN-dependent paracrine modulation mediated by TBK1 while TNF is unaffected

We next explored if the synergistic behaviors exhibited by these cytokines could be modulated by paracrine signaling. Since the dual combination of 100 ng/ml of P3C and 10 µg/ml of PIC consistently produced the highest levels of cytokine secretion for most targets and showed a synergistic effect for all (Fig. 2), we used this ligand combination for subsequent experiments. Previous studies in macrophages showed that in response to LPS, TNF and IFN-β can act in positive and negative feedback loops operating in a paracrine-dependent manner (Alexander et al., 2021; Gottschalk et al., 2019; Xue et al., 2015). We hypothesized that the synergistic responses observed in the TLR2-TLR3 combination are influenced by cell-cell communication through paracrine signaling pathways, including TNFR- and IFNAR-dependent pathways (Fig. 4A).

**Fig. 4.**
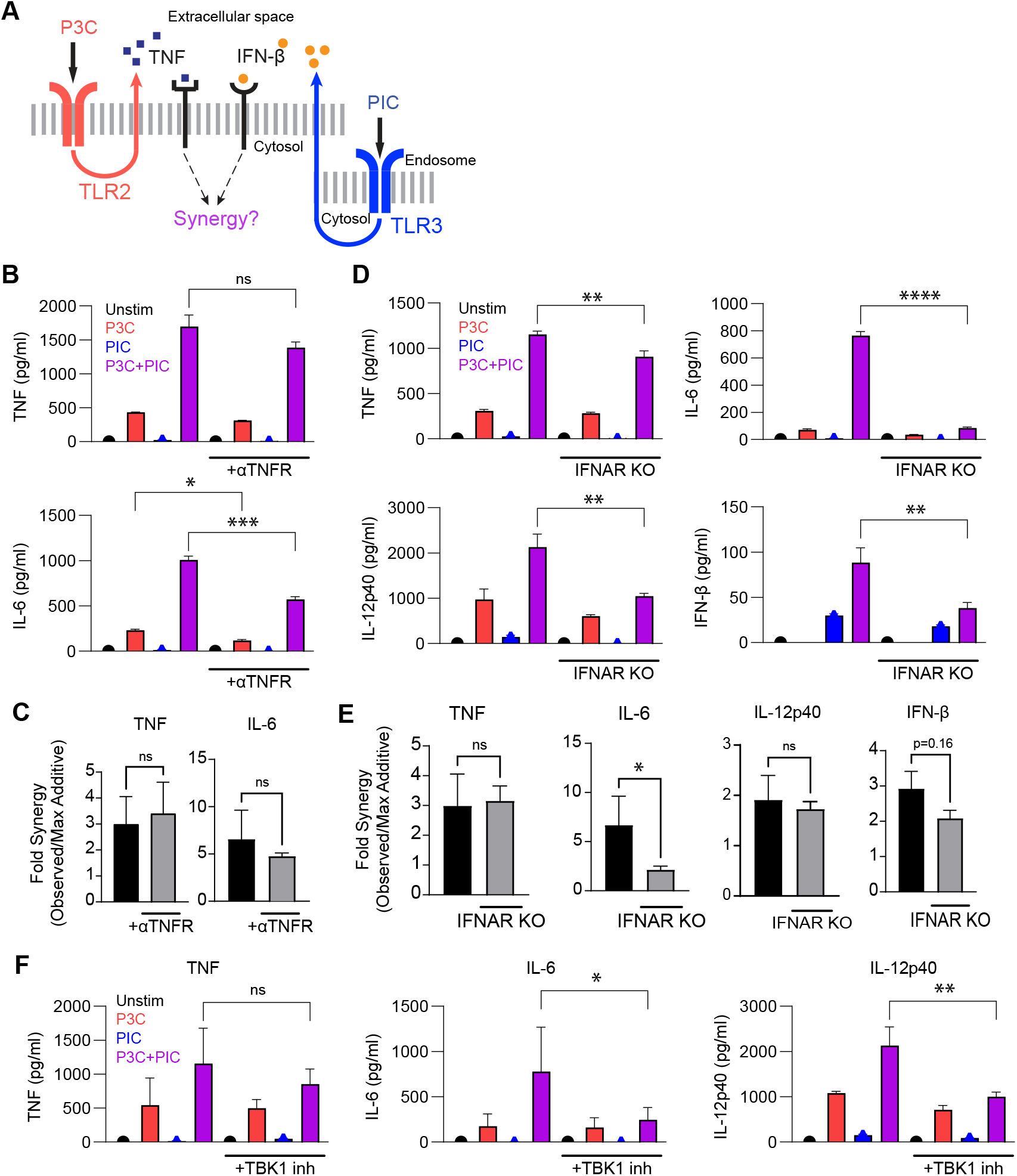
TLR2-TLR3 synergistic activation of IL-6 but not TNF exhibits type I IFN-dependent paracrine modulation mediated by TBK1. (A) Schematic of TLR2 and TLR3-stimulated paracrine signaling via TNFR and IFNAR. (B) WT BMDMs were stimulated with P3C and/or PIC alone and with anti-TNFR blocking Ab, and TNF and IL-6 secretion was measured by ELISA at 4 h post stimulation (n = 2). (C) Calculated fold synergy for the conditions in (B). (D) WT BMDMs or IFNAR KO BMDMs were stimulated with P3C and/or PIC, and cytokine secretion was measured at 4 h by ELISA for the indicated cytokines. (E) Calculated TLR2-TLR3 fold synergy between WT and IFNAR KO backgrounds. (F) BMDMs were stimulated with P3C and/or PIC +/-TBK1 inhibitor, and secretion was measured at 4 h by ELISA. Data for all bar graphs are presented as mean SD (n = 2). Significance was calculated using one-way ANOVA. Fold synergy was calculated between two experimental conditions and significance was determined by unpaired t-tests. ***p < 0.001, **p < 0.01, n.s. = not significant.

To determine if TNFR-mediated signaling contributes to TLR2-TLR3 synergy, we incubated BMDMs with a TNFR blocking antibody, challenged them with our ligand combinations. and measured the concentration of the pro-inflammatory cytokines TNF and IL-6 in the supernatants. TNF and IL-6 were determined to be the best targets from our panel to test, as they each include a primary and a secondary gene downstream of the NF-κB pathway (Liu et al., 2017). TNF secretion was unaffected by TNFR blockade, while IL-6 secretion decreased across the P3C and dual-ligand stimulations, consistent with a role for TNF in amplifying the IL-6 response that has been observed with LPS stimulation (Fig. 4B). However, in both cases, synergy was not significantly different (Fig. 4C). To confirm these observations, we stimulated BMDMs in the presence of a soluble TNFR (sTNFR) which would compete and reduce the availability of free TNF protein in the supernatants. As expected, we detected low levels of TNF in the supernatants (Fig. S4A) but we did not observe any significant difference for the secretion of IL-6 or its synergy (Fig. S4B). These results suggest that while TNF-signaling might contribute to the amplification of IL-6 secretion, it does not play a significant role in the synergy observed for either TNF or IL-6.

Type I interferons are key modulators of inflammatory responses through paracrine signaling (Michael Lavigne et al., 2021; Song et al., 2015). To assess a potential role for modulation of TLR2-TLR3 combination in a type I IFN-dependent manner, we used IFNAR KO BMDMs, challenged them with our ligand combinations, and measured the cytokine concentration of TNF, IL-6, IL-12p40, IFN-β, and CXCL10 in the supernatants. The lack of type I interferon signaling modestly but significantly reduced the production of TNF, IL-12p40, and IFN-β in the dual stimulation condition, while IL-6 secretion was more substantially reduced (Fig. 4D). As expected, we did not detect CXCL10 in the supernatants from IFNAR KO cells, consistent with their role as an ISG (Fig. S4C). Interestingly, IL-6 was the only target for which the loss of type I IFN signaling significantly reduced TLR2-TLR3 synergy (Fig. 4E). IFN-β showed a slight reduction in synergy, but it was not statistically significant (p = 0.16), which might be attributed to higher variability in the secretion of IFN-β in the WT BMDMs. Our combined results suggest that IL-6 synergy is modulated by paracrine signaling in an IFNAR-dependent manner.

Previous studies have shown that type I IFNs play an important role in tissue homeostasis and immune priming by tonic signaling (Gough et al., 2012; Ivashkiv & Donlin, 2014). We considered the possibility that constitutive levels of type I IFNs are needed for the synergy of IL-6 observed in the TLR2-TLR3 response, which would not be present in the IFNAR KO cells. To address that, we used WT BMDMs and intracellularly blocked the TRIF-dependent production of type I IFNs using a small-molecule inhibitor of TBK1 which is necessary for the downstream activation of IRF3 and the production of type I IFNs (Fitzgerald et al., 2003). We incubated these BMDMs one hour prior to stimulation with the TBK1 inhibitor, challenged them with our ligand combination of P3C and PIC, collected the supernatants at four hours, and measured cytokine concentration of TNF, IL-6, IL-12p40, IFN-β, and CXCL10 by ELISA. For the TBK1-inhibited cells, IFN-β and CXCL10 concentrations did not rise above the minimum level of detection by the assay (8 and 62.5 pg/mL, respectively), showing an effective block of type I IFN production (data not shown). The secretion of TNF and its synergy remained unaffected by TBK1 inhibition (Fig. 4F, left), suggesting that this branch of signaling is not necessary for the crosstalk that mediates the TNF synergistic response. In contrast, while TBK1 inhibition did not affect secretion of IL-6 in response to stimulation of P3C alone, IL-6 secretion in response to the dual stimulation was significantly decreased, with a complete inhibition of synergy (Fig. 4F, center). Lastly, IL-12p40 secretion did not show a significant decrease between the P3C stimulations, but the TBK1 inhibition resulted in a sharp decrease in synergy for the dual stimulation (Figure 4F, left) These results are consistent with those of our IFNAR KO cells, showing that IL-6 synergy requires type I IFN signaling.

### IFN-β alone is sufficient to modulate TLR2-mediated IL-6 and IL-12p40 secretion

We next investigated if murine IFN-β in combination with P3C could recapitulate the synergistic pro-inflammatory effect observed in the TLR2-TLR3 combination. We stimulated BMDMs with increasing doses of recombinant IFN-β protein (0, 0.01, 0.1, 1, 10, and 100 ng/mL) in combination with P3C. We observed that overall, compared to the P3C-only stimulation, IFN-β significantly increased P3C-mediated secretion of IL-6 in a dose-dependent manner (Fig. 5A, top). At the highest IFN-β dose of 100 ng/mL the synergy induced by IFN-β + P3C is indistinguishable from that of the P3C-PIC dual stimulation. Interestingly, when we measured IL-12p40 secretion, we observed that IFN-β increased P3C-mediated secretion only at the lower doses (0.01, 0.1, and 1 ng/mL), while at higher doses (1 and 10 ng/ml), IFN-β decreased P3C-stimulated secretion of Il-12p40, suggesting that IFN-β antagonizes rather than synergizes with TLR2 signaling at high doses (Fig. 5A, center). IFN-β did not significantly increase TNF secretion, and thus the synergy did not approach the level observed for PIC+P3C (Fig. S5A). We also tested how IFN-β affected secretion of IL-6 and IL-12p40 in combination with *S. aureus* infection (Fig. 5B). We found similar trends, with IFN-β increasing secretion of IL-6 at all doses but having a biphasic effect on IL-12p40 secretion, increasing secretion at low doses and inhibiting secretion at high doses.

**Fig. 5.**
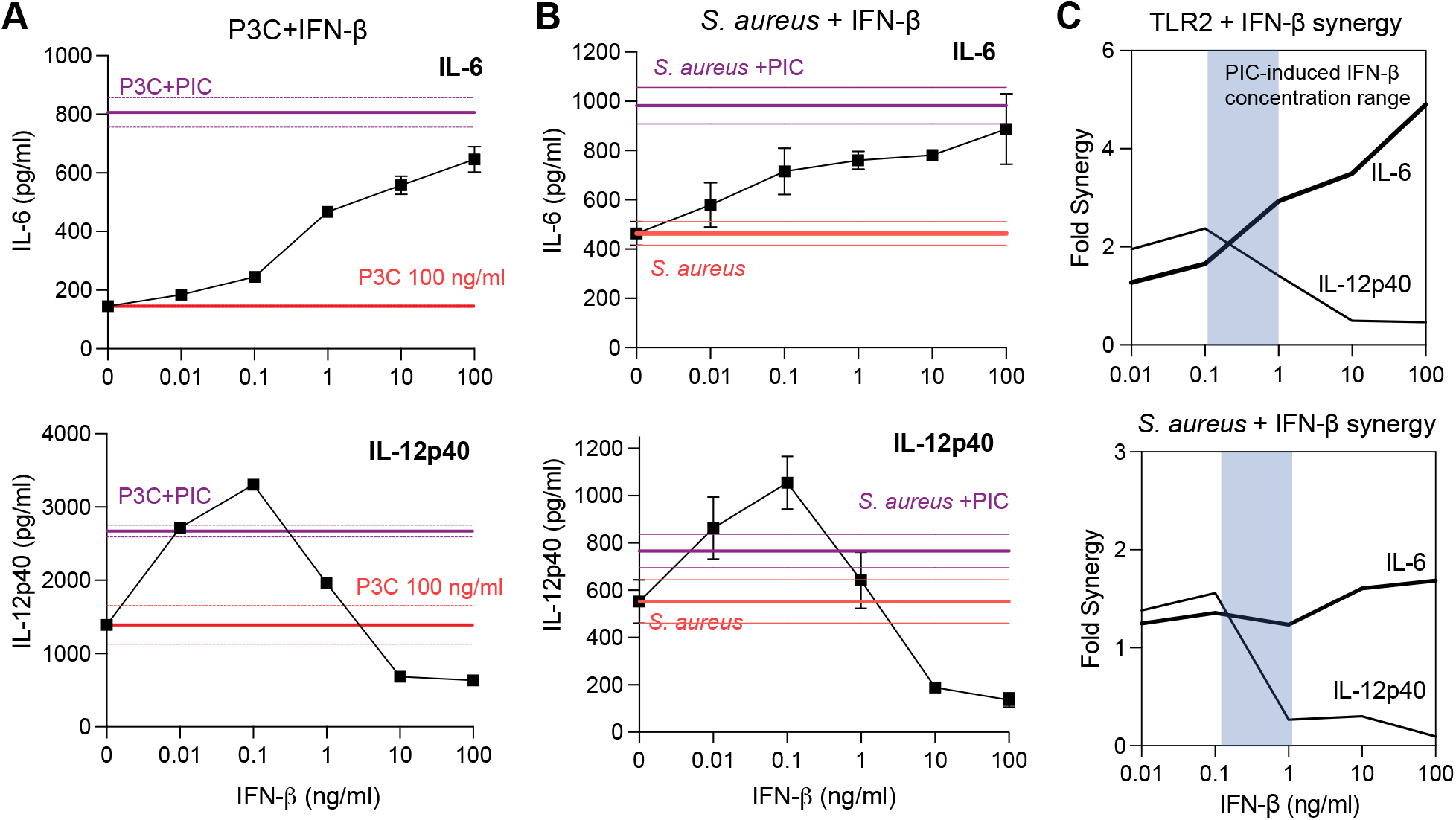
IFN-⊠ combines with TLR2 to induce synergistic activation of IL-6 but antagonizes activation of IL-12p40. (A) BMDMs were stimulated with 100 ng/ml of P3C and increasing doses of IFN-beta (0, 0.01, 0.1, 1, 10, 100 ng/ml) and the indicated cytokines were measured in the supernatants 4 hours post stimulation by ELISA. Reference lines (mean ⊠ SD) are presented for P3C only (red) and P3C+PIC (purple). (B) BMDMs were infected with S. aureus (MOI = 10) and increasing doses of IFN-β (0, 0.1, 1, 10, 100 ng/ml), and the indicated cytokines were measured in the supernatants 4 hours post infection by ELISA. Reference lines (mean SD) are presented for S. aureus infection only (red) and S. aureus + PIC (purple). (C) Fold Synergy of IL-6 and IL-12p40 was calculated for P3C or S. aureus in combination with increasing IFN-β doses. Data are presented as mean +/-SD (n = 2).

Calculating synergy as a function of IFN-β dose demonstrated the divergent trend between IL-6 and IL-12p40. Interestingly, in the range of IFN-β secretion induced by PIC, we found that synergy between IL-6 and IL-12p40 was relatively similar, but as the concentration of IFN-β increased, IL-6 synergy also continued to increase while IL-12p40 exhibited antagonism (Fig. 5C). These results highlight the role of IFN-β as an important paracrine modulator of both IL-6 and IL-12p40 TLR2-TLR3 synergy, but in different ways. At low concentrations of IFN-β, production of both IL-6 and IL-12p40 is synergistically increased. However, at high concentrations of IFN-β, production of IL-6 increases while the production of IL-12p40 decreases, demonstrating a potential trade-off in macrophages activating their interferon response.

### PIC and IFN-β both combine with other TLRs that signal through MyD88 to synergistically increase IL-6 secretion

Given that R848 stimulation of TLR7, which signals through MyD88, synergistically combined with PIC in a manner like TLR2, we next tested TLR5 and TLR9 stimulation with PIC. We stimulated BMDMs with flagellin (Flgn), a TLR5 ligand, and CpG oligodeoxynucleotides (CpG), a TLR9 ligand, each alone and in combination with PIC. As we observed for TLR2 and TLR7, co-stimulation with PIC synergistically upregulated IL-6 secretion (Fig. 6A). PIC also increased IL-12p40 secretion with modest synergy when combined with TLR7 and TLR5 but not for TLR9. However, PIC did synergistically activate TNF when combined with TLR9 (Fig. S5B). This finding suggests that, while the mechanism for PIC-mediated synergy changes for some cytokines, IL-6 synergy is similar across all TLRs that activate MyD88.

**Fig. 6.**
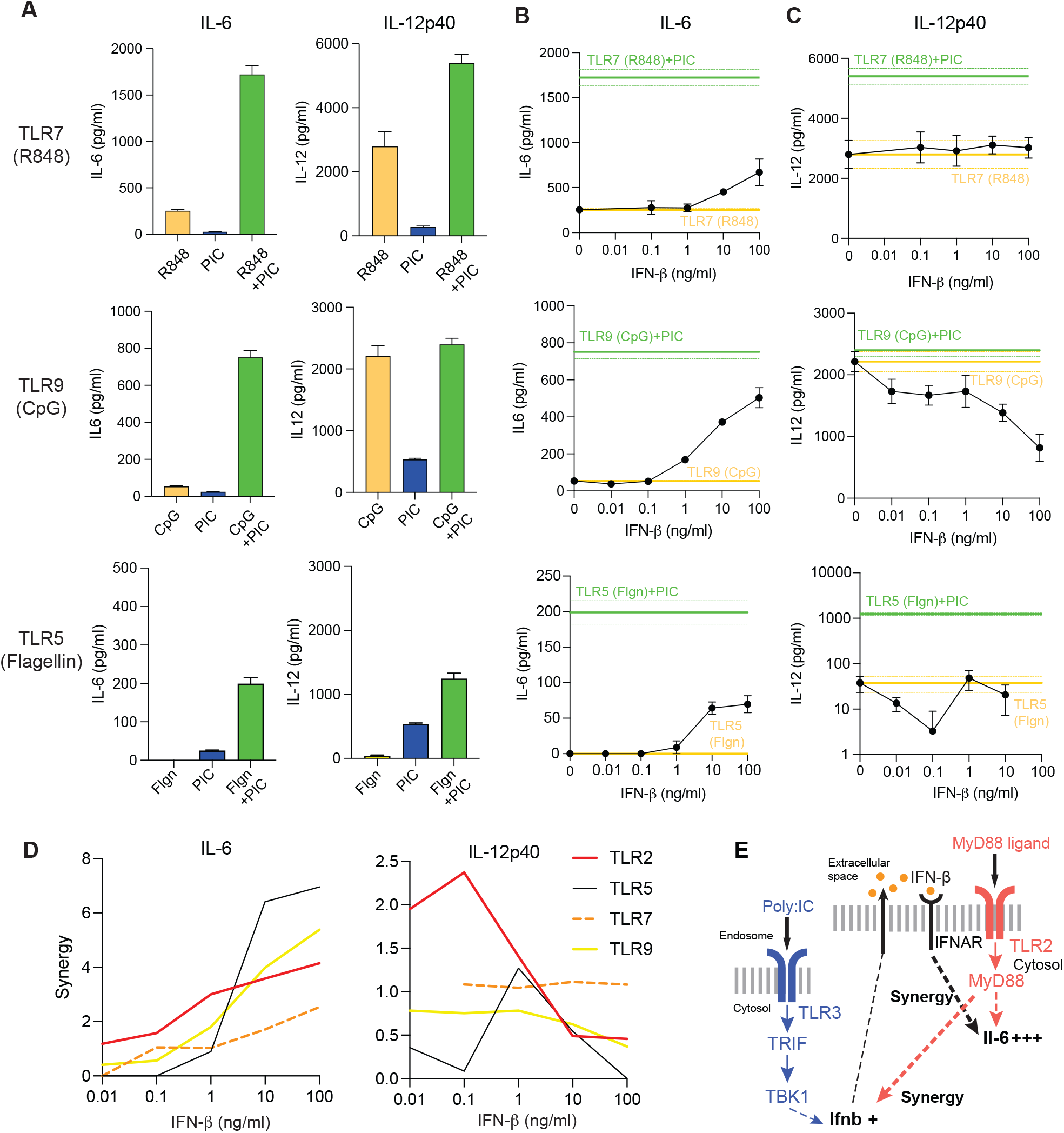
PIC and IFN-β induce synergistic activation of IL-6 across a range of TLR ligands that signal via MyD88. (A) BMDMs were stimulated with 10 µg/ml PIC and 100 ng/ml R848, 100 ng/ml flagellin, 1 µM CpG, or the combination for 4 hours and then IL-6 and IL-12p40 secretion was measured by ELISA. (B) BMDMs were stimulated with the same doses of the MyD88 ligands as in (A) and increasing doses of IFN-beta (0, 0.01, 0.1, 1, 10, 100 ng/ml) and IL-6 (B) and IL-12p40 (C) were measured in the supernatants 4 hours post stimulation by ELISA. Reference lines (mean +/-SD) are presented for TLRX only (yellow) and TLRX+PIC (green). (D) Fold Synergy of IL-6 and IL-12p40 was calculated for each TLR ligand with increasing IFN-β doses. Data are presented as mean SD of n = 2 biological replicates. (E) Schematic of the mechanism of IL-6 synergy mediated by IFN-β.

We then tested how IFN-β alone modified IL-6 and IL-12p40 secretion when combined with other TLRs that signal through MyD88. As before, we varied IFN-β from 0.01 to 100 ng/ml in combination with each TLR ligand. We found that IFN-β increased IL-6 secretion in combination with each TLR at doses at or above 1 ng/ml (Fig. 6B). In contrast, the effect of IFN-β on IL-12p40 secretion was varied from neutral to antagonism at high doses of IFN-β for TLR9 (Fig. 6C). IFN-β only modestly increased activation of TNF by TLR9 stimulation at the highest doses (Fig. S5C). When we compared all MyD88-activating TLR ligands, we found that IFN-β exhibited a clear trend of increased synergy with increasing dose (Fig. 6D). In contrast, IFN-β variably modulated secretion of IL-12p40 induced by MyD88-activating TLR ligands at low doses but generally antagonized IL-12p40 secretion at high doses. Overall, we conclude that IFN-β can directly increase activation of the Th1 cytokines IL-6 and IL-12p40, and that this is an important mechanism by which TLR3 synergizes with MyD88-activating TLR ligands to amplify IL-6 production.

## Discussion

The TRIF and MyD88 pathways mediate responses to different members of the TLR family. There has been a long-standing interest in how TLR2, mediated by MyD88, and TLR3, mediated by TRIF, combine to activate the innate immune response in macrophages. Here we showed that co-stimulating TLR2 and TLR3 in murine macrophages with P3C and PIC, respectively, synergistically activates cytokines involved in the inflammatory and antiviral response. Synergistic activation of secretion was higher than for transcription, and it varied by time and ligand dose. Importantly, our results support a new mechanism for IL-6 synergy observed when combining TLR3-TRIF ligands with ligands that signal via MyD88 pathways (Fig. 6E). IFN-β secretion is activated in response to TLR3 stimulation–and can be further amplified via MyD88-dependent pathways that have yet to be elucidated. IFN-β can then specifically amplify secretion of IL-6 in response to TLR-MyD88 ligands, accounting for at least part of the synergy observed.

Several earlier studies explored mechanisms of synergy between TLR2 and TLR3 in macrophages and other cell types. Synergistic effects are observed in TLR combinations that engage both the MyD88 and TRIF signaling pathways regardless of cellular localization (surface vs endosomal) or the specific ligand (*e*.*g*., lipoproteins, LPS, viral nucleic acids, etc.) (Bagchi et al., 2007; Ouyang et al., 2007). While no definitive mechanism has been elucidated for the synergy of MyD88-TRIF, it requires *de novo* protein synthesis and a complex interplay between transcription factors from the MAPK, C/EBP, AP-1, and IRF transcription factor families (Liu et al., 2015; Suet Ting Tan et al., 2013). Moreover, previous studies have suggested that Type I IFNs might play a role in synergy, as evidenced by studies in BMDMs showing enrichment of IRF and STAT transcription factors in MyD88/TRIF combinatorial stimulations that are not observed in single-ligand stimulations, and studies in human DCs showing a modest role in IL-12p40 secretion (Gautier et al., 2005; Lin et al., 2017). Despite this observation, IFN-β and other antiviral genes have been poorly characterized in the context of multi-TLR combinations. Furthermore, although previous studies have noted that IL-6 is synergistically activated by TLR2 or TLR7 (MyD88) in combination with TLR3 (TRIF), no study has directly implicated IFN-β as a major mediator of the synergy.

While IFN-β alone does not stimulate the production of IL-6 or IL-12, it has been shown to modulate TLR-mediated IL-6 and IL-12p40 production in other contexts. There is evidence that type I interferons increase the expression and production of IL-6. For example, IFN-α increased IL-6 production in response to TLR8 stimulation in human neutrophils (Zimmermann et al., 2016). There is evidence that type I interferons inhibit IL-12 production in mouse splenic leukocytes and in dendritic cells, as well as *in vivo* during viral infection (Cousens et al., 1997; McRae et al., 1998). IFN-β also causes a dose-related inhibition of IL-12p40 and p70 production by primary human monocytes (Karp et al., 2000). Although it has been suggested that IFN-β inhibition of IL-12 might be mediated by IL-10, these studies showed that it was largely independent of IL-10.

There is a growing appreciation that immune responses can be amplified via paracrine signaling from a fraction of the population (Antonioli et al., 2019). In the case of IFN-β, it was observed that production is stochastic, with only a fraction of infected cells producing IFN-β (Rand et al., 2012; Zhao et al., 2012). However, this fraction of cells activates ISGs in neighboring cells via paracrine signaling to establish a protective antiviral state (Patil et al., 2015; Rand et al., 2012). It is interesting to consider if there is a biological advantage to further coupling amplification of IL-6 secretion to anti-viral protection. One possibility is that it would allow the surviving cells to produce sufficient Th1 cytokines to make up for those cells that succumbed to viral infection. However, this would need to be tested in an in vivo challenge to dual TLR stimulation.

Our findings contribute to a growing body of evidence that type I interferons (IFN-I) can act as amplifiers of IL-6 secretion during combinatorial TLR signaling, suggesting a mechanism that may help explain how immune responses escalate in both protective and pathological contexts. This has relevance for diseases characterized by IFN-I dysregulation, such as STING-associated vasculopathy with onset in infancy (SAVI)—a monogenic autoinflammatory disease caused by constitutive STING activation—as well as severe COVID-19. In SAVI, chronic overproduction of IFN-I leads to sustained IL-6 secretion and systemic inflammation, while in severe COVID-19, a delayed but exaggerated IFN-I response correlates with IL-6-driven cytokine storms and tissue damage (Ramasamy & Subbian, 2021; Wang et al., 2021). An IFN-β–IL-6 amplification loop suggests that even limited IFN-producing cells may initiate a broader paracrine cascade with significant consequences for inflammation. By clarifying how type I IFNs regulate cytokine synergy, this work supports the development of precision immunotherapies aimed at either promoting or restraining cytokine responses in disease-specific contexts.

## Methods

### Mouse breeding and cell culture

Mice were maintained in specific pathogen-free conditions in the Yale Animal Resources Center (YARC) and all procedures were performed according to the approved protocols of the Yale University Institutional Animal Care and Use Committee. Female C57BL/6J (wild-type) and *Ifnar1*^*-/-*^(IFNAR KO) mice 6-8 weeks of age were purchased from Jackson Laboratories. Mice were housed at room temperature according to the standard housing conditions of YARC. Bone marrow-derived macrophages (BMDMs) were generated as previously described (Trouplin et al., 2013). Briefly, bone marrow was extracted from the tibias and femurs of the hind legs of the mouse with a syringe. After red blood cell lysis with ammonium-chloride-potassium lysis buffer (Lonza), cells were incubated for 4 h at 37 °C with 5% CO_2_ in a non-tissue culture treated plastic petri dish with BMDM media (RPMI supplemented with 10% FBS, 100 U/ml penicillin, 100 μg/ml streptomycin, 1% sodium pyruvate, 25 mM HEPES buffer, 2 mM L-glutamine, and 50 μM 2-mercaptoethanol). After 4 h, the non-adherent cells were transferred to a new petri dish and incubated with BMDM media supplemented with 20 ng/ml macrophage-colony stimulating factor (M-CSF; Peprotech). Three days after plating, an additional 10 ml of BMDM media supplemented with 20 ng/ml M-CSF was added to the plate. On day 6, cells were rinsed with PBS and lifted in PBS with 5 mM EDTA through gentle pipetting, and cells were plated in non-TC treated 24-well plates at a density of 250,000 cells/well and were allowed to adhere overnight.

### *In vitro* BMDM treatments

BMDMs were plated and stimulated in BMDM media with 10 ng/ml M-CSF (PeproTech). Cells were stimulated with Pam3CSK4 (Invivogen), poly(I:C) HMW (Invivogen), recombinant murine IFN-β (R&D), ultrapure flagellin from *S. typhimurium* (Invivogen), CpG ODN 1668 (Invivogen), and R848 (Invivogen) alone, in combination, or co-stimulated with a TNFR1 blocking antibody (Invitrogen) clone 55R-170 or a small-molecule TBK1 inhibitor (GSK8612; MedChemExpress) for the time and doses indicated in the figure legends. Cells used for intracellular cytokine staining or blocked for paracrine communication were treated with Golgi Plug™ (BD Biosciences), a Brefeldin A (BFA)-containing compound for the duration of the incubation (four hours).

### Bacterial strains and infections

Bacterial strains used were the following: *Staphylococcus aureus subsp. aureus NCIMB 12702* (Microbiologics) and an isogenic *dotA flaA* mutant *Legionella pneumophila* in the LP02 background, a streptomycin-resistant thymidine auxotroph derived from *L. pneumophila* LP01, with in-frame deletions of the *dotA* and *flaA* genes (Ren et al., 2006). Lyophilized *S. aureus* was re-hydrated and grown on a Trypticase Soy Agar petri dish according to Microbiologics protocol and incubated at 37 °C until colony formation. The day prior to infection, single colonies were grown in Trypticase Soy Broth overnight (12-16 hours), and absorbance at 600 nm was measured to determine final bacterial concentration. Bacteria were washed two times in cold PBS and seeded BMDMs were infected at an MOI of 10 by centrifugation for 5 min at 200 x g. *Legionella* strains were grown on charcoal yeast extract plates (1% yeast extract, 1% *N*-(2-acetamido)-2-amino-ethanesulphonic acid (pH 6.9), 3.3 mM L-cysteine, 0.33 mM Fe(NO_3_)_3_, 1.5% bacto-agar, 0.2% activated charcoal) supplemented with 100 µg/ml thymidine. *Legionella* were grown for 2 days at 37 °C, harvested, and absorbance at 600 nm was measured to determine final bacterial concentration. For all *Legionella* experiments, BMDMs were infected at an MOI of 10 by centrifugation for 5 min at 200 x g.

### RT-qPCR

RT-qPCR was performed as previously described (Wasik et al., 2016). In brief, cells were lysed, and RNA was extracted using the RNEasy Mini Kit (Qiagen). Genomic DNA was removed on-column with RNase-free DNase (Qiagen), and complementary DNA (cDNA) was synthesized using a dT oligo primer and Superscript III RT (Invitrogen). cDNA was eluted in nuclease-free water and was quantified using SYBR-green for quantitative polymerase chain reaction on a CFX Connect Real-Time System (Bio-Rad) with the following amplification scheme: 95 °C denaturation for 1.5 min followed by 40 cycles of 95 °C denaturation for 10 s, 65 °C annealing for 10 s, and 72 °C elongation for 45 s with a fluorescence read at the end of the elongation step. This was followed by a 65–85 °C melt-curve analysis with 0.5 °C increments. All samples were normalized to the house-keeping gene *Ubc* (Ubiquitin C). Primers are listed in Table 1.

**Table 1.**
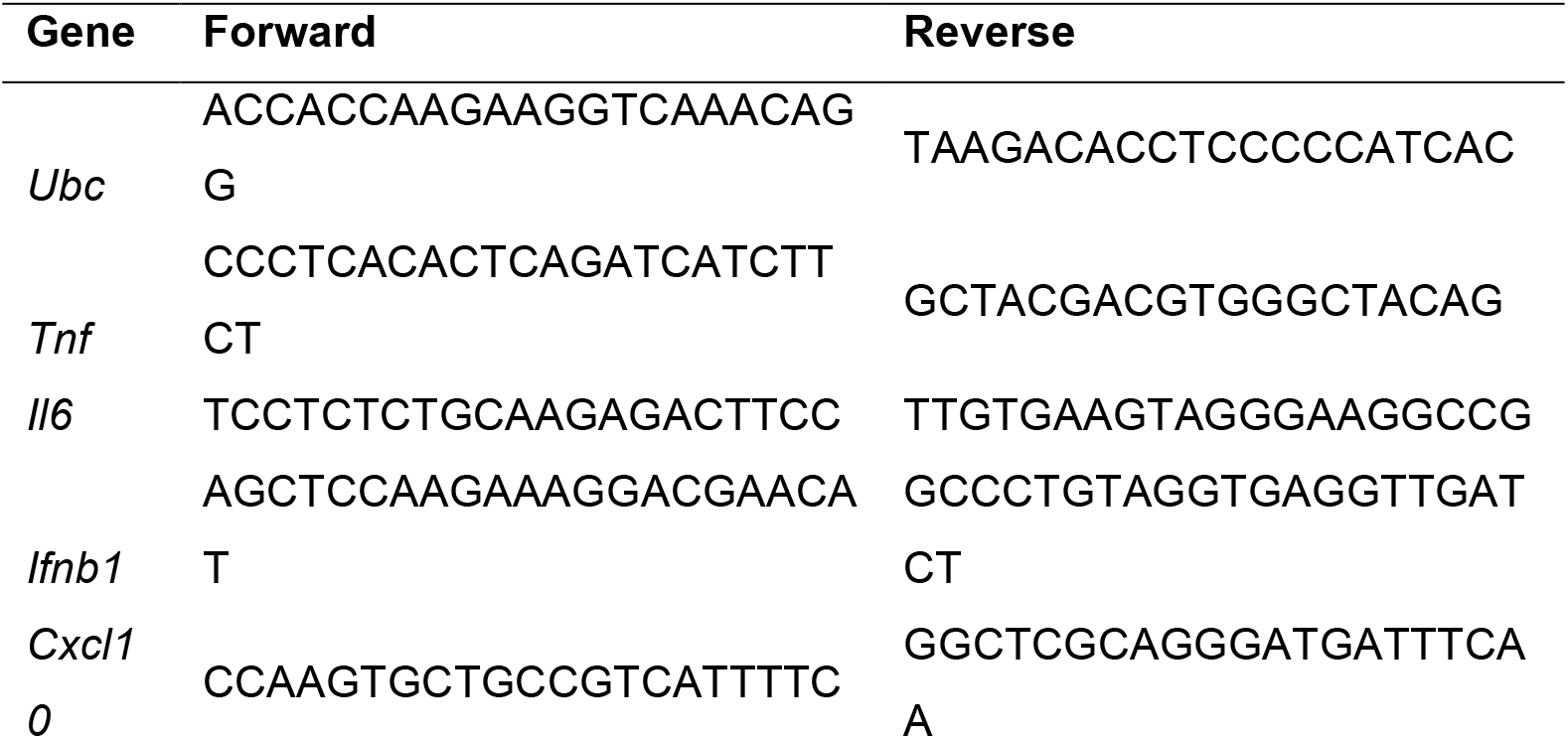
List of primer sequences.

### Secretion measurements

Cell culture media from stimulated BMDMs was collected at the end of the incubation period. Secreted protein levels were measured using enzyme-linked immunosorbent assay (ELISA) kits according to the manufacturer’s instructions. Each ELISA kit reference number and manufacturer are listed in Table 2. For multiplex secretion measurements 100 µL of cell culture media were collected and then submitted to Eve Technologies (Calgary, AB, Canada) to perform a Mouse Cytokine/Chemokine Array 32-Plex (MD32) assay, which is based on color-coded polystyrene beads combined with a dual-laser and a flow-cytometry system for sample acquisition and analysis.

**Table 2.** List of capture and detection antibody pairs.

| Antibody Pair | Vendor | Catalog No. |
| --- | --- | --- |
| TNF | Invitrogen | 88-7324-88 |
| IL-6 | R&D | DY406 |
| IL-12p40 | BD Biosciences | BDB555165 |
| IFN-β | R&D | DY823405 |
| CXCL10 | R&D | DY46605 |

### Flow Cytometry

Macrophages were lifted by gentle pipetting in ice-cold PBS with 5 mM EDTA. Cells were incubated with FcBlock (anti-CD16/CD32, eBiosciences) at 1:200 dilution on ice for 15 min in FACS buffer (PBS 2% FBS). Cells were then washed and fixed with Cytofix/Cytoperm and Perm/Wash Buffer kit (BD Biosciences) according to manufacturer’s instructions and stained in 100 μl with BV421 anti-mouse TNF-alpha (MP6-XT22) and PE-Cy7 anti-mouse IL-12p40 (C15.6) at a 1:100 dilution for 30 min at 4 °C. All data were acquired on an LSRFortessa (BD Biosciences), and analyzed with FlowJo (FlowJo, LLC). The gating strategy is shown in Supplementary Fig. 2.

## Statistics

Data were presented as mean ± SD unless otherwise specified. Statistical analysis was generally performed by two-sided, unpaired Student’s *t*-test or one-way ANOVA and the Sidak method of correction for pairwise multiple comparisons. Normal and equal distribution of variances was assumed. Values were considered significant at *P* < 0.05. All analyses were performed using Prism version 10.4.1 software (GraphPad). Clustering of multiplex secretion data was performed using the ‘clustergram’ function in MATLAB (Mathworks). Measured supernatant concentrations were Log2 transformed before being mean-centered and variance scaled for each protein.

## Notes

### Competing Interest Statement

The authors have declared no competing interest.

### Funding

- National Institutes of Health: R01GM123011, T32AI055403

