## Supplemental Figures for "Interferon-β paracrine signaling mediates synergy between TLR3 and TRIF-independent TLR pathways"

Supplemental Material

Supplemental Figures 1-5

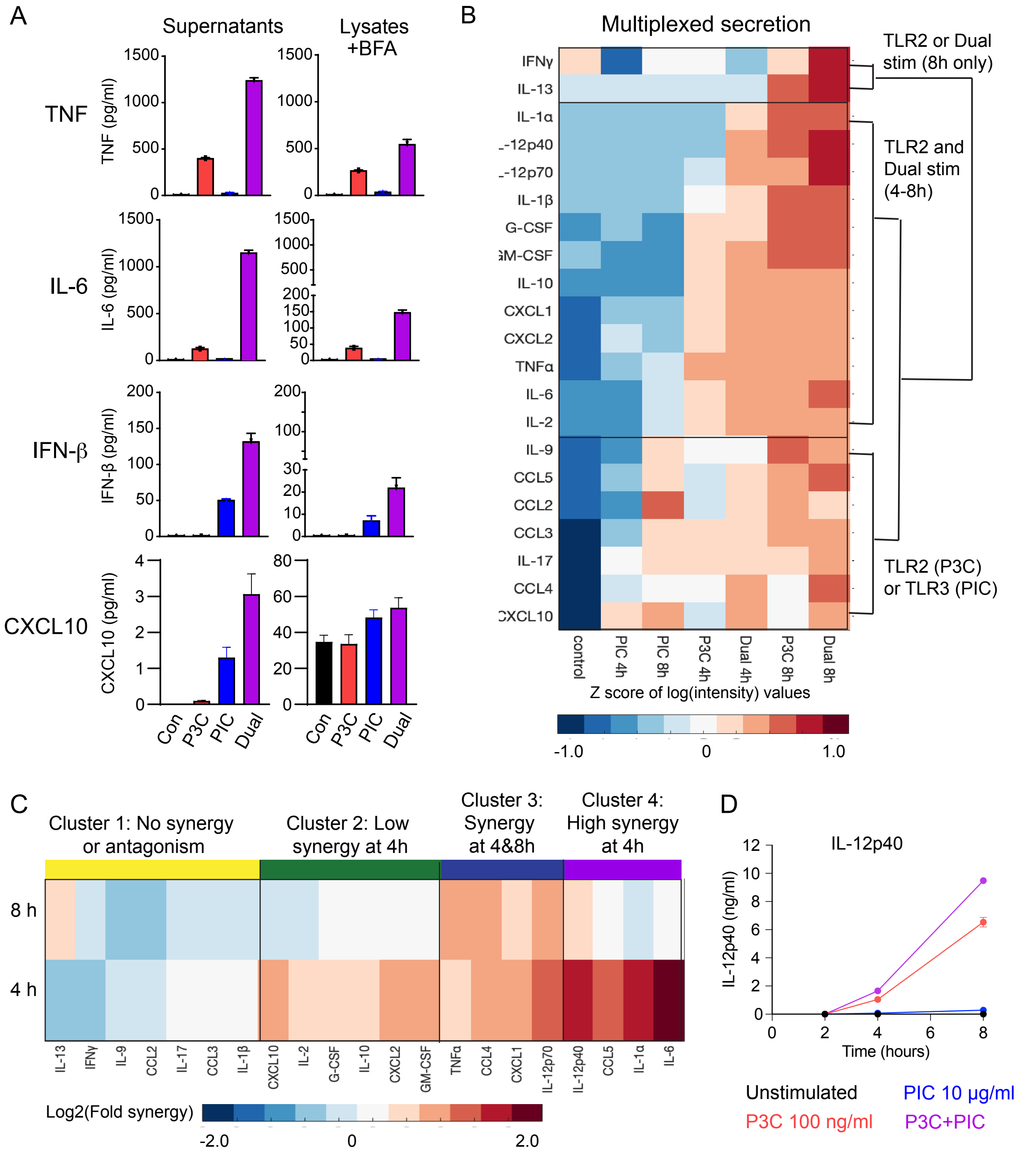


**Fig. S1. TLR2-TLR3 synergy in protein production is observed widely across C/Cs.** (A) BMDMs were treated with P3C, PIC, and P3C+PIC in combination with Brefeldin A, the BMDMs were lysed four hours post stimulation, and the indicated cytokines were measured in these lysates (n = 2). (B-C) Secretion was measured in the supernatants of BMDMs stimulated with P3C, PIC, and P3C+PIC at 4 and 8 hours, and multiplexed secretion was measured by bead-based immunoassay. (B) Cytokines and chemokines that were significantly activated above control were Z-scored and grouped by hierarchical clustering. (C) Fold synergy was calculated for each cytokine and grouped based on hierarchical clustering. Multiplexed data, n = 2 (control), n = 1 (treatments). (D) Results from multiplexed data in (C) were confirmed by ELISA for IL-12p40. Data are presented as mean   SD of n = 3 biological replicates.

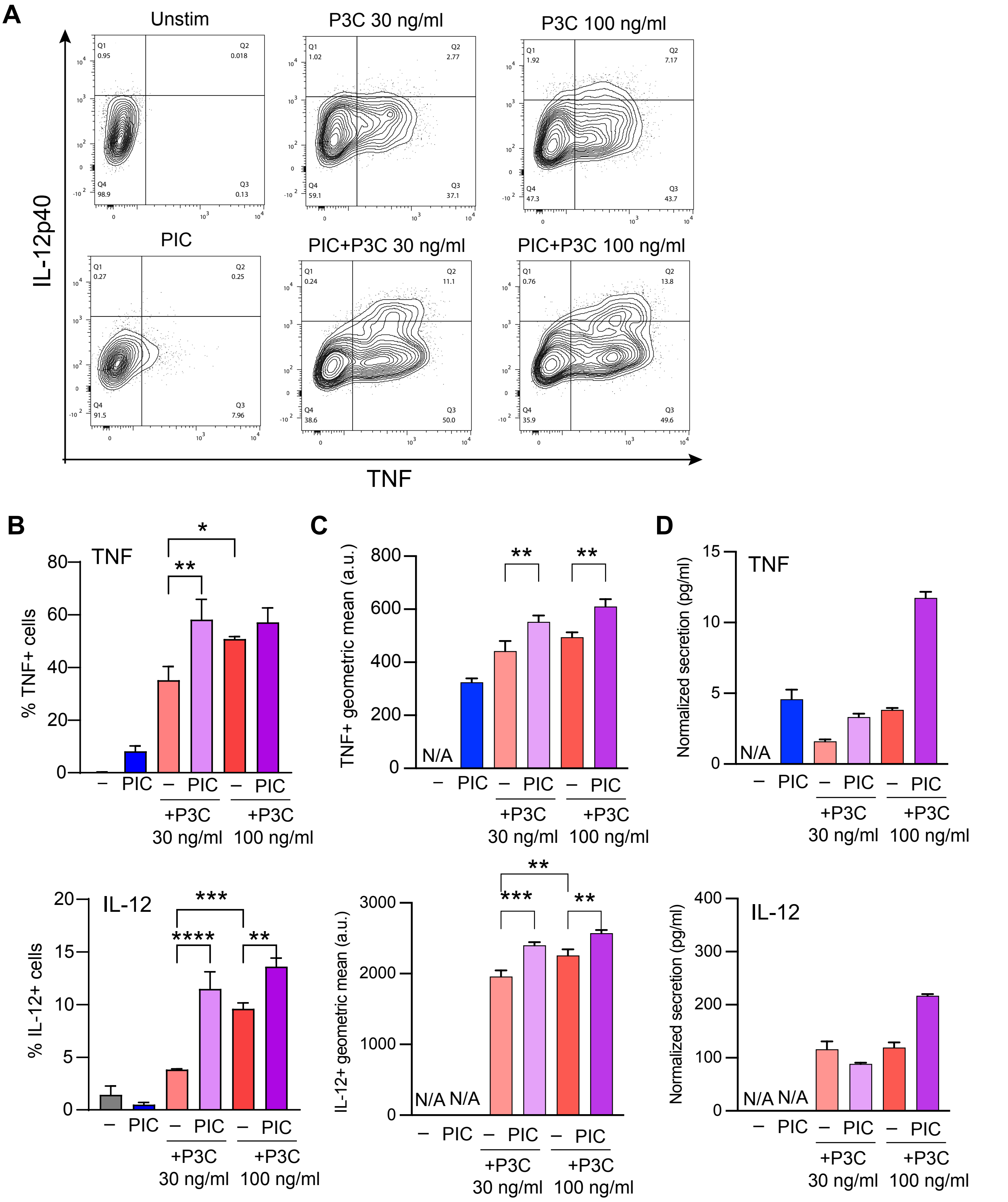


**Fig. S2. Synergistic activation of TNF and IL-12p40 increases both the fraction of cells producing each cytokine and the per cell amount.** (A) Sample flow plots and gates for TNF+ and IL-12p40+ cells. (B) Percent positive cells for TNF (top) and IL-12p40 (bottom). (C) Geometric mean of the positive fraction. (D) Total secretion (as measured in Fig. 2) normalized by the fraction of cells activated. Data are presented as mean SD of n = 2 biological replicates.

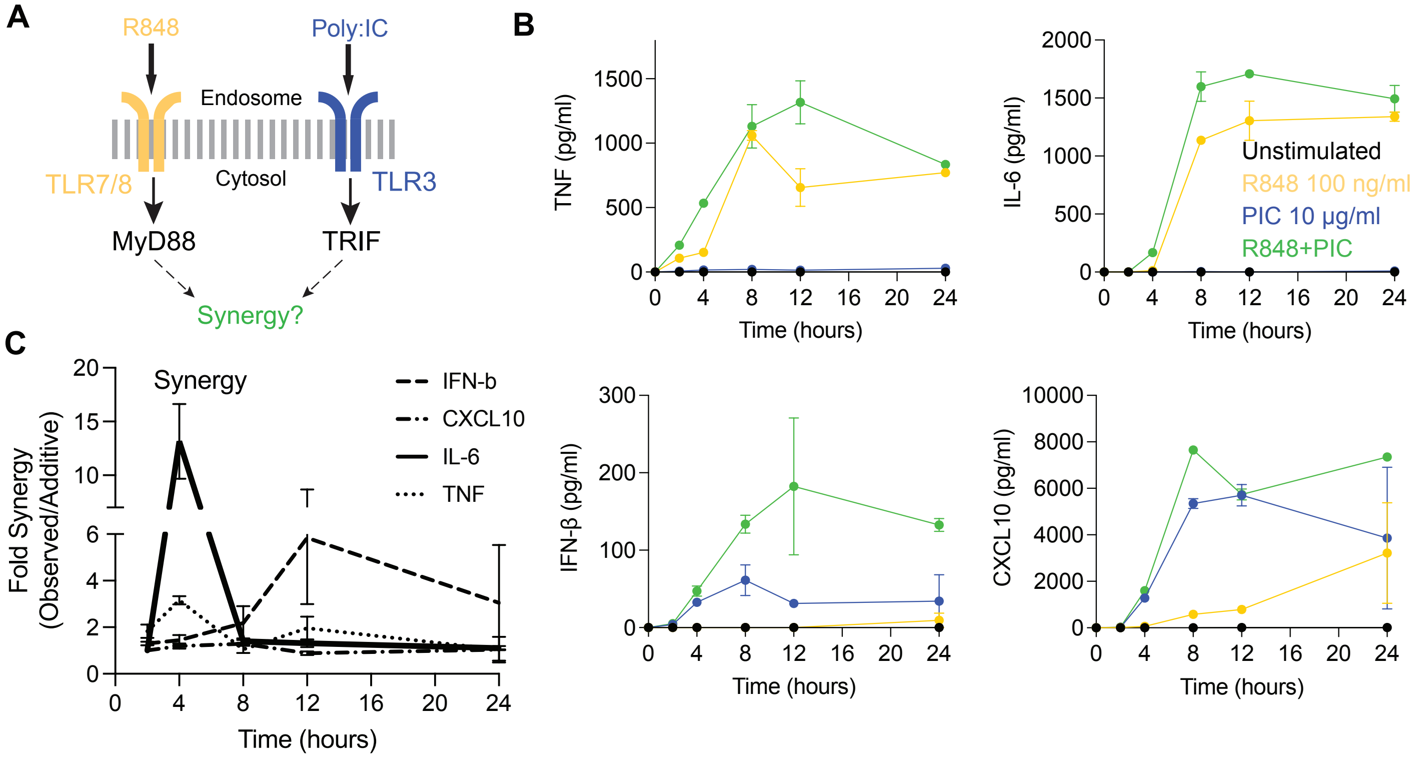


**Fig. S3. TLR7/8 stimulation with R848 also synergizes with PIC-TLR3 stimulation.** (A) Schematic of TLR pathways. (B) ELISA measurements of protein concentrations of indicated targets in the supernatants of BMDMs stimulated with R848 and/or PIC at 2 to 24 hours. (C) Calculated fold synergy of the R848+PIC secretion response for all measured cytokines across timepoints. Fold synergy = response to R848+PIC/(response to R848 + response to PIC). All graphs present data as mean SD of n = 2 biological replicates.

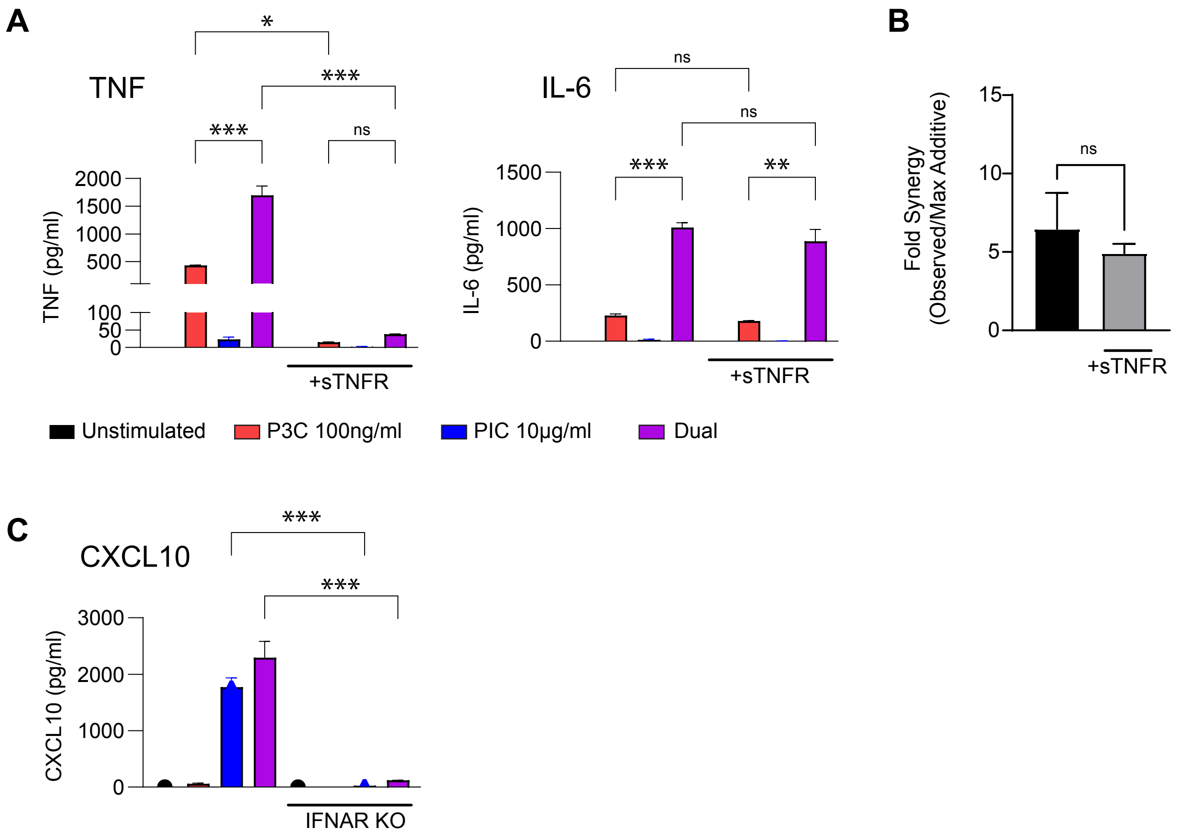


**Fig. S4. Blocking paracrine signals reveals role for type I interferons but not TNF in mediating synergy.** (A) BMDMs were treated with P3C (100 ng/ml), PIC (10 µg/ml), and the combination for 4 hours with and without soluble TNFR to block TNF paracrine signaling. TNF and IL-6 were measured by ELISA. Data represent the mean SD of n = 2 biological replicates. (B) Calculation of the change in synergy for IL-6. (C) Experiment as in (A) but with BMDMs from IFNAR knockout mice. CXCL10 measured by ELISA.

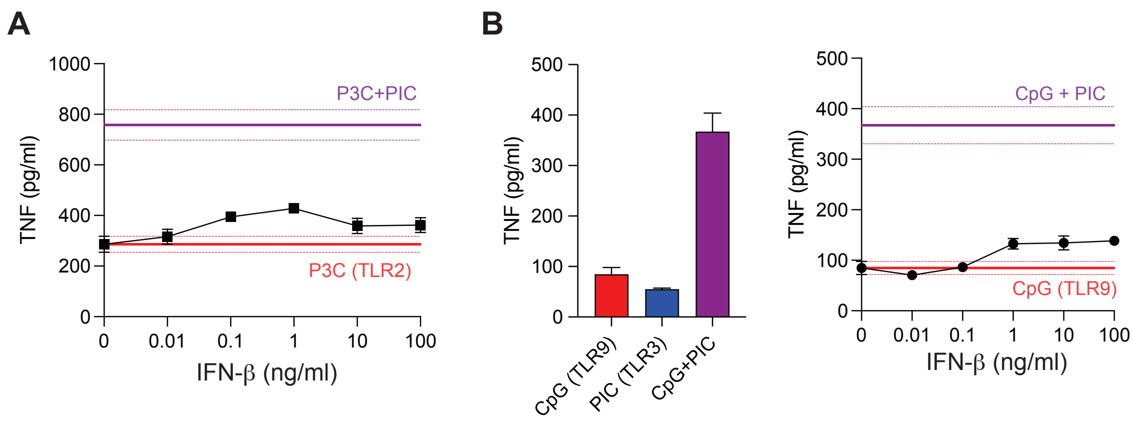


**Fig. S5. IFN-β does not synergistically activate TNF upon co-stimulation with TLR2 or TLR9 ligands.** (A) BMDMs were stimulated with 100 ng/ml of P3C and increasing doses of IFN- (0, 0.01, 0.1, 1, 10, 100 ng/ml) and TNF was measured in the supernatants 4 hours post stimulation by ELISA. Reference lines (mean +/- SD) are presented for P3C only (red) and P3C+PIC (purple). (B-C) BMDMs were stimulated with 10 µg/ml PIC and 1 µM CpG (B) or increasing doses of IFN-β and TNF was measured in the supernatants 4 hours post infection by ELISA. Reference lines (mean +/- SD) are presented for CpG only (red) and PIC + CpG (purple). Data presented as mean SD of n = 2 biological replicates.
